# Quantitative Real-Time Imaging of Glutathione with Sub-Cellular Resolution

**DOI:** 10.1101/360362

**Authors:** Xiqian Jiang, Chengwei Zhang, Jianwei Chen, Sungwoo Choi, Ying Zhou, Mingkun Zhao, Xianzhou Song, Xi Chen, Mirjana Maletić-Savatić, Timothy Palzkill, David Moore, Meng C. Wang, Jin Wang

## Abstract

Quantitative imaging of glutathione with high spatial and temporal resolution is essential for studying the roles of glutathione in redox biology. We developed a fluorescent glutathione probe—HaloRT—that targets organelles of interest through expressing organelle-specific HaloTag proteins. Using HaloRT, we quantitatively measure the glutathione concentrations in the nucleus and the cytosol and find no appreciable concentration gradient between these two organelles, challenging the view of nuclear compartmentalization of glutathione.

## Main text

Glutathione (GSH), the most abundant non-protein small peptide in eukaryotic cells, regulates the intracellular redox homeostasis with its oxidized disulfide partner, GSSG. Imbalanced redox states have been shown to lead to various abnormal conditions such as diseases and drug resistance^1,2^. Understanding the intracellular concentration and distribution of GSH provides the most direct evidence to localized cellular redox state and is the key step to study GSH-related redox biology. Just as pH is essential to all cellular events, GSH level is also crucial to a considerable number of biochemical processes inside cells. Importantly, the activity of GSH is tightly linked to its spatial and temporal dynamics, which however remains poorly characterized due to a lack of effectives method for quantifying subcellular GSH levels in real-time.

Our group pioneered the use of reversible reaction-based ratiometric fluorescent probes for measuring GSH concentrations in live cells. We developed the first fluorescent probe ThiolQuant Green (TQG) that can quantitatively measure intracellular GSH levels in live cells^3^. Building on this work, we developed the 2^nd^ generation GSH probe RealThiol (RT) which allows monitoring dynamic changes of GSH levels in real time^4^. The groups of Urano and Yoon also independently reported two probes that can follow GSH dynamics ^5,6^. Recently, we developed a mitochondria-targeted GSH probe, MitoRT ^7^, taking advantage of the high binding affinity of a triphenylphosphonium moiety towards the highly negatively charged mitochondrial membrane ^8^ and revealed the specificity of mitochondrial GSH dynamics. Unfortunately, this small molecule-based subcellular targeting strategy is not suitable to all different organelles. To develop a universal strategy that can be easily and broadly applied to any organelles of interest, we designed a HaloTag enabled GSH probe, named HaloRT (Fig.1). Here, using different signaling peptide sequences, we are able to specifically locate HaloRT in either the nucleus or the cytosol, and measure organelle-specific GSH concentrations. To our surprise, we discovered that there is no appreciable GSH concentration gradient between the nucleus and the cytosol in HeLa cells and in primary hepatocytes under several different treatment conditions, which strongly challenges the existing view of a nuclear pool of GSH ^9^.

**Fig.1.**
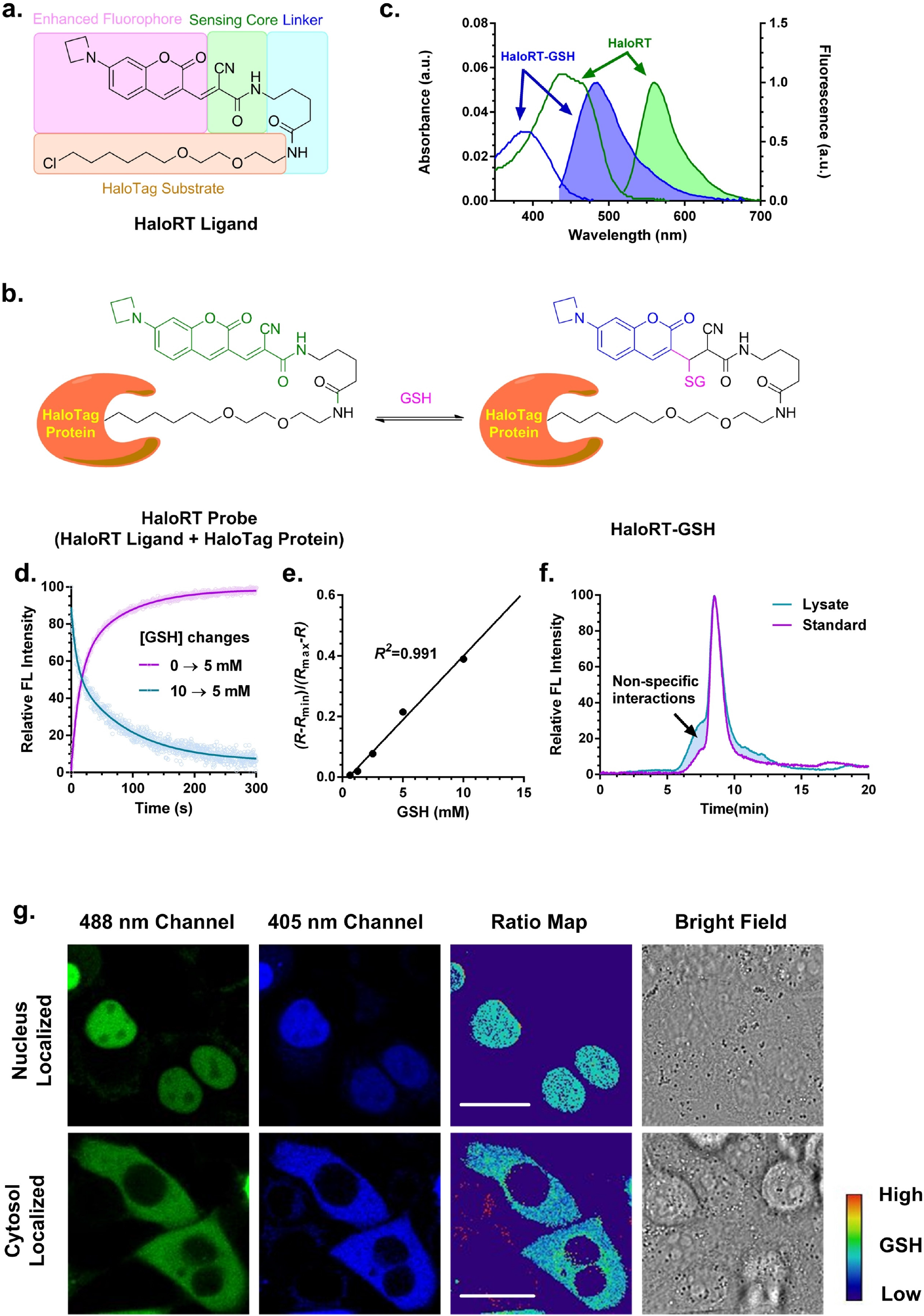
Characterization of HaloRT probe. (**a**) Chemical structure of HaloRT ligand and the modular design diagram. (**b**) Reversible reaction between HaloRT and GSH. (**c**) Deconvoluted UV-Vis absorption spectra (unshaded) and fluorescent spectra (shaded) of HaloRT (Green, *λ*_ex_ = 488 nm) and HaloRT-GSH conjugate (blue, *λ*_ex_ = 405 nm). (**d**) Reaction kinetics of HaloRT and GSH measured on a stopped-flow rapid mixing device with a fluorescence detector. Forward and backward reactions were monitored at *λ*_ex_ = 405 nm, *λ*_em_ = 480 nm, which correspond to the formation and dissociation of HaloRT-GSH conjugate. (**e**) Calibration curve determined by mixing a series of known concentrations of GSH with HaloRT followed by imaging on a coverslip using confocal microscopy. The calculated *K*_d_’ is 23.7 mM, and the *R*^2^ is 0.991 in the physiological relevant concentration range of GSH. (**f**) Gel permeation chromatography trace of standard HaloRT-GSH conjugate and lysate from cyto-HaloRT stained HeLa cells. Fluorescent signal (*λ*_ex_ = 405 nm, *λ*_em_ = 480 nm) was used for detecting HaloRT-GSH conjugate. Calculation of the area under the curve revealed that 87% of the signal can be attributed to HaloRT-GSH, indicating high selectivity towards GSH over protein thiols inside cells. (**g**) Representative confocal images of nuclear and cytosolic specific HaloRT stained HeLa cells. Ratio maps were calculated by dividing the blue channel by the green channel at each corresponding pixel. Scale bars are 10 µm.

To achieve sub-cellular specificity for GSH probes, we adopted the widely-used HaloTag technology for organelle specific protein tagging^10^. Derived from our GSH RT probe, we substituted the di-carboxylic acid moiety with the HaloTag protein substrate elongated with a four-carbon linker (HaloRT ligand, Fig. 1a, supplementary note 1 and Fig. S1). Then, by expressing organelle-specific HaloTag proteins in live cells, the HaloRT ligand can be recruited to different organelles and used for quantitative real-time imaging of GSH at sub-cellular resolution.

After successfully synthesizing the HaloRT ligand, we first verified that HaloRT, the conjugate between the HaloRT ligand and recombinant HaloTag protein, exhibits similar GSH sensing abilities as RT. Similar to RT, HaloRT exhibits a bright orange color and an absorption maximum at 480 nm, and shows fluorescence maximum at 565 nm upon excitation at 488 nm. Importantly, addition of GSH shifts the absorption and fluorescence peaks to 390 and 488 nm (excited at 405 nm), respectively (Fig. 1b and 1c).

The kinetics between HaloRT and GSH is also very similar to that of RT. We measured the reaction using a stopped-flow rapid mixing device at room temperature and monitored the formation and disappearance of HaloRT-GSH by fluorescence (λ_ex_ = 405 nm, λ_em_ = 480 nm). We observed that both forward and reverse reactions equilibrate in about 200 s (Fig. 1d). To measure the dissociation equilibrium constant (*K*_d_) of the reversible sensing reaction, we mixed a series of known concentrations of GSH (from 0.39 to 50 mM) with HaloRT (~10 µM) and measured the fluorescence in two channels (λ_ex1_ = 405 nm, λ_em1_ = 480 nm; λ_ex2_ = 488 nm, λ_em2_ = 565 nm). The *K*_d_ for the sensing reaction is 121 mM, 32 folds higher than that of RT. The low reactivity of the protein conjugated RT may be due to the steric hindrance of the protein or the hydrophobic HaloTag protein surface shifts the *K*_d_ of thiol-Michael addition reactions. It should be noted that the apparent dissociation equilibrium constant (*K*_d_’), which is dependent on instrument settings, is more relevant in determining the dynamic range of the probe. By adjusting the gains of both emission channels, *K*_d_’ values are 24.8 and 23.7 mM measured by a plate reader and a confocal microscope used in this study. In both cases, the calibration curve confers a superb linear relationship with *R*^2^ above 0.99 within the physiological relevant GSH concentration range (1.25-15 mM, Fig. 1e and Fig. S2). The high *K*_d_ of HaloRT results in a very small percentage of HaloRT-GSH formed under physiological conditions. Compared to RT, we increased the gain for the 405 nm excitation channel to achieve sensitive detection of the subtle changes of HaloRT-GSH level.

Next, we determined if the probe is selective towards GSH in cells, and applied the gel permeation chromatography (GPC) method with fluorescence detection that we developed for assessing the selectivity of RT in complicated intracellular environment ^4^. The lysate of HaloRT stained cells were washed using a denature buffer at pH 4.5 to prevent reverse reactions, and the fluorescence signal of HaloRT-GSH (λ_ex_ = 405 nm, λ_em_ = 480 nm) was used for detection. Compared to the HaloRT-GSH standard underwent the same processing, we did not observe significant molecular weight changes (Fig. 1f), suggesting that the HaloRT probe selectively reacts with small molecule thiols but not protein thiols in cells. Together, based on the fluorescence spectra, kinetic and thermodynamic constants, and the selectivity profile, we conclude that HaloRT is suitable to quantitatively monitor GSH level changes under cellular conditions in real-time.

To express the HaloTag protein in different organelles, we fused it with organelle targeting sequences, Halo-NLS with a nuclear localization sequence and Halo-MapKK with a cytosolic localization sequence from MapK kinase (MapKK) ^11^. HeLa cells were transiently transfected with Halo-NLS and Halo-MapKK, incubated with the HaloRT ligand at 37 °C for 30 minutes, and then imaged using a confocal microscope. We confirmed that Halo-NLS and Halo-MapKK specifically target the HaloRT ligand to the nucleus (nuc-HaloRT) and the cytosol (cyto-HaloRT), respectively (Fig. 1g). Thus, our strategy is effective to target the GSH probe to different cellular organelles.

It is generally accepted that GSH is compartmentalized in different organelles and regulated separately there, but whether there is a nuclear pool of GSH remains a long-standing debate ^12-14^. Bellomo et al. was the first to report a nuclear pool of GSH in primary hepatocytes using monochlorobimane (MCB) as a GSH probe ^12^. However, this study was challenged by Briviba et al. based on their observation that MCB-GSH conjugate microinjected into the cytosol can preferentially accumulate in the nucleus, suggesting that MCB could react with cytosolic GSH first and the conjugate is further enriched in the nucleus ^13^. Although some other studies favor the view of nuclear pool of GSH, the fluorescent probes used suffer the same artefact as MCB ^9^. In our system, the readouts of HaloRT are independent of probe concentrations due to its ratiometric nature. The Halo-NLS and Halo-MapKK are first transfected into the cells and expressed into their targeted organelles before seeing the HaloRT ligand. Additionally, we have confirmed that nuc-HaloRT and cyto-HaloRT have high spatial specificity. Thus, we have applied our system to reexamine the nuclear GSH theory. We discovered no appreciable concentration gradient of GSH between the nucleus and the cytosol (supplementary note 2). Despite the absolute fluorescence signals from the nucleus and cytosol are slightly different due to the variation of the transfection efficiencies, the ratios of fluorescence signals remain statistically the same in these two locations. This ratiometric feature of our GSH probes is a unique advantage to avoid the non-specific influence from different transfection efficiencies and probe availability (Fig. 2b and supplement note 2).

**Fig.2.**
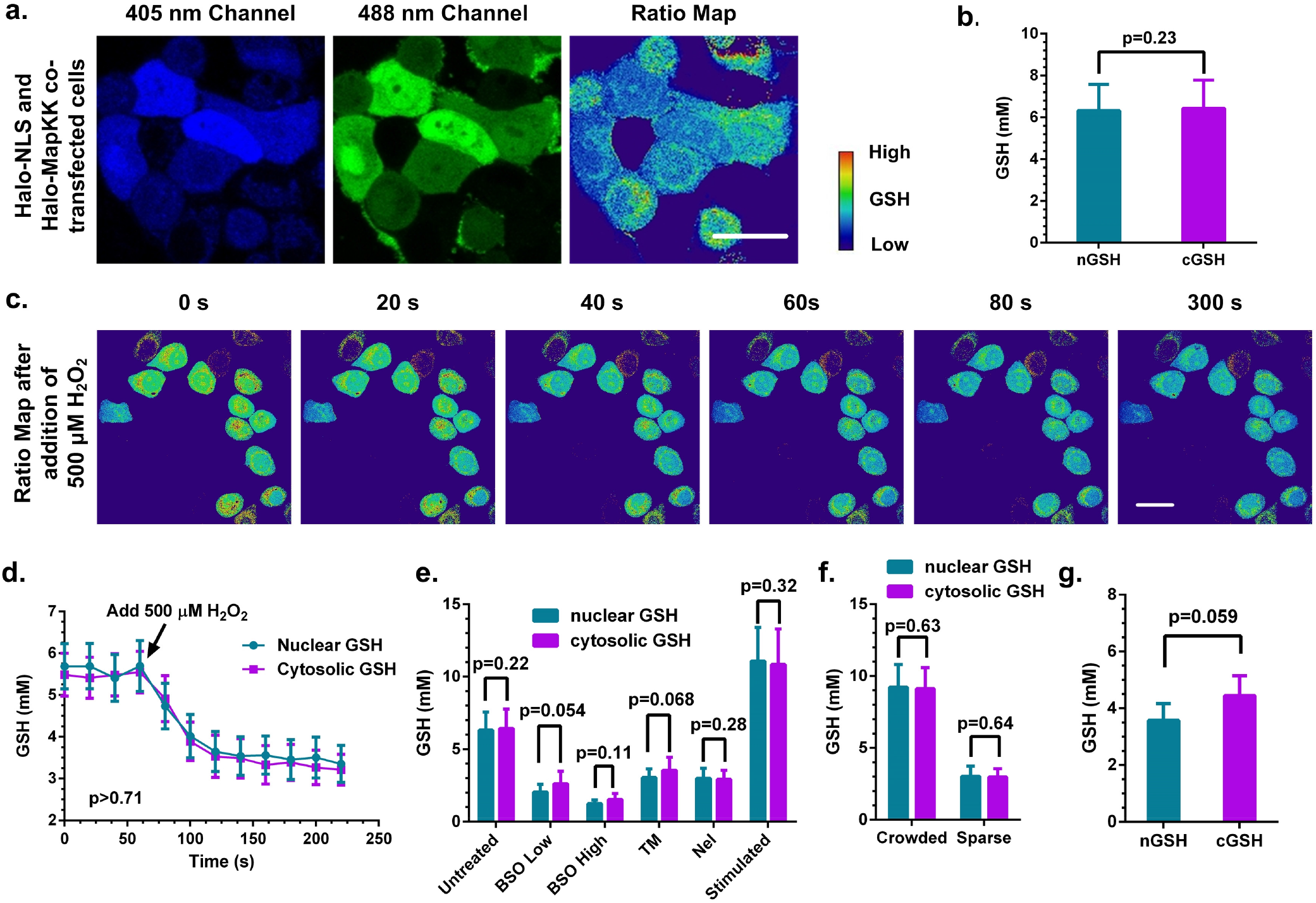
Organelle-specific HaloRT probes show no nuclear pool of GSH. (**a**) Representative images of HaloRT ligand stained HeLa cells co-transfected with both nuclear specific and cytosolic specific HaloTag protein. Scale bar is 10 µm. (**b**) Quantitative analysis of GSH concentrations in the nucleus and the cytosol from untreated HeLa cells. A total of 394 and 420 cells from >25 images from >5 individual experiments were statistically analyzed for nGSH and cGSH groups, respectively. (**c**) Time-lapsed ratio maps from HeLa cells transiently treated with 500 μM of H_2_O_2_ at time 0. Scale bar is 10 µm. (**d**) Quantitative analysis of GSH concentrations from transient H_2_O_2_ treated HeLa cells. A total of 15 and 18 cells from the representative experiment group show in (**c**) were statistically analyzed for nGSH and cGSH groups, respectively. (**e**) Quantitative analysis of GSH concentration from HeLa cells treated with vehicle, 50 μM of buthionine sulfoximine (BSO) for 24 h, 500 μM of BSO for 24 h, 10 μM of tunicamycin (TM) for 4 h, 10 μM of Nelfinavir (Nel) for 4 h, and 100 μM of diethyl-maleate (DEM) for 2h then allowed to recover for 22h, respectively (from left to right). A total of 394, 188, 229, 178, 172, and 184 cells from >25 images from >5 individual experiments were statistically analyzed for nGSH group at each corresponding treatment conditions, respectively; and a total of 420, 219, 262, 234, 236, and 198 cells from >25 images from >5 individual experiments were statistically analyzed for cGSH group at each corresponding treatment conditions, respectively. (**f**) Quantitative analysis of GSH concentrations from HeLa cells cultured in crowded region and in sparse region. A total of 122 and 65 cells from >15 images from >3 individual experiments were statistically analyzed for nGSH group in the two regions, respectively; and a total of 119 and 96 cells from >15 images from >3 individual experiments were statistically analyzed for cGSH group in the two regions, respectively. (**g**) Quantitative analysis of GSH concentrations from untreated primary hepatocytes. A total of 38 and 46 cells from >15 images from >3 individual experiments were statistically analyzed for nGSH and cGSH groups, respectively. Standard unpaired t test was used for all statistical analyses. All the error bars represent standard deviations.

Next, to investigate whether the nuclear GSH (nGSH) can be better maintained compared to cytosolic GSH (cGSH), we treated Halo-NLS and Halo-MapKK co-transfected HeLa cells with different compounds and measured the corresponding nGSH and cGSH under these conditions. In a time-lapsed experiment, we treated cells with a bolus of 500 μM of H_2_O_2_ to induce oxidative stress, and imaged for 5 min consecutively. As shown in Fig. 2c, both nGSH and cGSH levels decrease synchronously but there is no significant GSH gradient between the nucleus and the cytosol throughout this period. Further quantification demonstrates that nGSH and cGSH levels are not different from each other before and after H_2_O_2_ treatments (Fig. 2d).

We also challenged cells with other stressors and examined their effects on nGSH and cGSH. Buthionine sulfoximine (BSO) is a specific GSH synthase inhibitor which decreases intracellular GSH in a concentration dependent manner. When we treated HeLa cells with 50 or 500 μM of BSO for 48 h, we found that both nGSH and cGSH decreases as BSO concentration increases (Fig. 2e and Fig. S3), but their levels are statistically the same. Endoplasmic reticulum (ER) is a critical cellular compartment for protein folding, modification and quality control, which is linked to the oxidizing balance of GSH. We thus have treated cells with tunicamycin (TM) and nelfinavir (NEL) to induce ER stress^15^ and test whether these treatments would affect nGSH and/or cGSH levels. We found that after 4 h of treatments, nGSH and cGSH are decreased to similar levels (Fig. 2e), suggesting that ER stress-induced oxidative insults affect both the nucleus and the cytosol. Together, these results show that shifting the oxidative balance toward oxidation reduces nGSH and cGSH levels at the same time and to the same degree, which do not support an independent pool of GSH in the nucleus.

On the other hand, we examined whether an increase in GSH would affect the concentration gradient between nGSH and cGSH. To this end, we stimulated cells with 100 μM of diethyl maleate (DEM) transiently for 2 h and then allowed the cells to recover for 24 h. We found that this treatment greatly increases GSH levels in the cell, but there is no difference between nGSH and cGSH increases (Fig. 2e and Fig. S3). Densely cultured cells are known to have higher GSH levels compared to sparsely cultured counterparts ^16,17^. We thus generated a cell culture condition where crowded and sparse regions co-exist within the same culture dish and show 2-3 folds of GSH concentration difference (Fig. S4). Under this physiological condition without exogenous stimulants, we did not observe an appreciable difference between nGSH and cGSH (Fig. 2f). Furthermore, we also examined nGSH and cGSH levels in primary hepatocytes transfected with Halo-NLS and Halo-MapKK and stained with the HaloRT ligand. Consistent with our observation in cell line studies, there is no appreciable difference between nGSH and cGSH in these primary cells (Fig. 2g and Fig. S5). Therefore, no concentration difference between nGSH and cGSH with increased GSH levels and under physiological conditions.

In conclusion, we have developed an organelle-specific GSH probe, HaloRT, which can be universally applied to determine GSH levels in any organelles of interest with targeted HaloTag protein expression. This HaloRT system enables us to quantify GSH concentrations with sub-cellular resolution in real time. Taking advantage of this new system, we revisited the nuclear GSH theory. Our measurements do not reveal any appreciable GSH concentration gradients between the cytosol and the nucleus and thus do not support an independent pool of nuclear GSH. Considering the nuclear pore complex has an inner diameter of approximately 9-12 nm and typically allows free diffusion of molecules less than 60 kDa ^18,19^, the small tripeptide GSH should freely diffuse through the nuclear pores. Although demonstrated only for the nucleus and the cytosol, with the HaloRT probe in hand, this system can be applied to any organelles, and will be a valuable addition to the redox biology toolbox.

## Methods

### Chemical synthesis of HaloRT ligand

Refer to the supplementary note 3 for details.

### Synthesis of HaloRT probe

HaloRT ligand (400 μM, 200 μL) was incubated with purified His-HaloTag protein (Promega, G4491, 6.5 mg/mL, 50μL) for 1 h at room temperature in PBS. The reaction mixture was then concentrated with a 10k cut-off viva-spin 500 centrifugation tube, and washed twice with fresh PBS. The resulting bright orange supernatant was then used as HaloRT probe for further testing. The HaloRT probe was validated by gel permeation chromatography to ensure that no free HaloRT ligand remained in solution.

### UV-vis and fluorescence spectra of HaloRT probe

HaloRT probe was diluted with PBS to 10 μM and mixed with equal volume of PBS or GSH (700 mM). The resulting mixture was then measured on a BioTek H1 plate-reader for absorption and fluorescence at corresponding wavelengths using a Take3 Trio micro-volume plates. A deconvolution of the overlapping spectra was performed to yield the individual spectrum.

### Kinetics of HaloRT probe reacting with GSH

The reaction kinetics was measured on a stopped-flow rapid mixing device with a fluorescence detector. The formation and dissociation of HaloRT-GSH conjugate was monitored at *λ*_ex_ = 405 nm, *λ*_em_ = 480 nm. The forward reaction was measured by mixing equal volume of 10 μM HaloRT probe with 10 mM of GSH and the reverse reaction was measured by diluting a pre-equilibrated 10 μM HaloRT probe and 10 mM GSH mixture with an equal volume of PBS.

### Equilibrium constant of the reaction between HaloRT probe and GSH

HaloRT probe was diluted with PBS to 10 μM and mixed with equal volume of a series of known concentrations of GSH (0.39, 0.78, 1.56, 3.13, 6.25, 12.5, 25, 50, 100 mM) and equilibrated for 5 min at room temperature. The resulting mixture was then measured on plate-reader at two fluorescent channels: *λ*_ex_ = 405 nm, *λ*_em_ = 480 nm and *λ*_ex_ = 488 nm, *λ*_em_ = 565 nm. The ratio between the two channels was used for calibration of equilibrium constant.

Similarly, the mixture of HaloRT probe and GSH was prepared and imaged using a confocal microscope. Before imaging, the mixture was mixed with 15 μm polystyrene beads for focusing purpose. A typical sample can be prepared by sealing 5 μL of solution with a coverslip.

### Intracellular selectivity of HaloRT probe measured with GPC-FL

To acquire enough amount of material for analysis, we choose to over-express HaloTag protein in cytosol which has the highest overall quantity of protein. HeLa cells (from a confluent 15 mm dish) transiently transfected with cytosolic specific HaloTag protein were stained with 5 μM of HaloRT ligand at 37 °C for 30 min. The cells were washed twice with cold PBS to remove excessive small molecule probes, and then lysed using 1 mL of 10% trichloroacetic acid on ice. The lysate was then centrifuged at 15,000 g at 4 °C for 15 min. The supernatant was ready for GPC-FL measurement as the small molecule mixture. The resulting precipitate was then redissolved using 0.1 M pH 4.5 citric acid buffered 5% SDS at 58 °C for 5 min. Supernatant was then collected after quickly centrifuging at 15,000 g at room temperature for 1 min. This second supernatant was ready for GPC-FL measurement as the re-dissolved protein mixture.

GPC-FL measurement was performed using 0.1 M pH 4.5 citric acid buffered 0.1% SDS solution with a cationic column. Flow rate was set at 1.5 mL/min and the temperature was set at 45 °C. A standard HaloRT-GSH sample, a small molecule mixture from cell lysate, and a re-dissolved protein mixture from the same cell lysate were measured on GPC-FL with fluorescent detector set at *λ*_ex_ = 405 nm, *λ*_em_ = 480 nm for detection of HaloRT-GSH conjugate. The small molecule mixture resulted in no fluorescent signal, and the HaloRT-GSH standard and the redissolved protein mixture showed similar retention time on their fluorescent signal. Area under the curve (AUC) after normalization was used for quantification purpose. Analysis revealed that 87% of the protein mixture signal comes from HaloRT-GSH as compared with the standard.

### General cell culture

The HeLa cell line used in this study was purchased from American Type Culture Collection (ATCC) and grown in DMEM (Gibco, 11965) media supplemented with 10% FBS and 1% 1003 Pen Strep (Gibco). Cells were cultured under a controlled atmosphere (37 °C, 5% CO_2_). Glass bottom dishes were used for cell culture due to confocal scanning requirements.

### Organelle specific quantitative imaging of GSH using HaloRT probe

Halo-NLS and Halo-MapKK plasmids were constructed by integrating organelle specific signal sequences into Halo cDNA sequence. The following signal sequences are used for plasmid constructs: 5’-CCAAAAAAGAAGAGAAAGGTAGAAGACCCC-3’ for nuclear specific, 5’-TTGCAGAAGAAGCTGGAGGAGCTAGAGCTTGAT-3’ for cytoplasm specific. Whole piece of DNA fragment, including signal sequence and Halo cDNA sequence, was synthesized by Genscript, and then inserted into pCDNA 3.1 (+) vector through KpnI (5’end) and NotI (3’end) sites.

HeLa cells (4×10^5^) were seeded on a 35-mm glass bottom culture dish and incubated for 24 h. The cells were then transfected with one or two plasmids using Lipofectamine 3000 transfection agent and incubated for additional 36 h. Before imaging, cells were stained with 1 μM HaloRT ligand in culture medium at 37 °C for 30 min, and then washed with fresh medium twice, each for 10 min. Imaging was performed using a Zeiss LSM 780 confocal microscope with a temperature control chamber. Green channel signal was acquired with λ_ex_ = 488 nm, λ_em_ = 499-615 nm; blue channel signal was acquired with λ_ex_ = 405 nm, λ_em_ = 418-495 nm.

### Treatment conditions for HeLa cells

Transient treatment with H_2_O_2_ was performed at the site of imaging. Stained, transfected HeLa cells were treated with a bolus of 500 μM of H_2_O_2_ (100 μL 5 mM H_2_O_2_ in PBS diluted to 1 mL medium) in the imaging chamber and imaged at 37 °C continuously for 5 min with 20 s per scan.

BSO (50 μM and 500 μM) was added to HeLa cells at seeding, and the cells were incubated for 24 h before transfection of cells.

DEM (100 μM) was added to HeLa cells 10 h after transfection. Cells were incubated with DEM for 2 h and then washed with fresh culture medium. Then, the cells were incubated for additional 22 h in fresh culture medium before imaging.

ER stressors (TM 10 μM and Nel 10 μM) were added to cells 4 h prior to imaging.

HeLa cells were seeded on a confined spot on a culture dish without stirring, which created a crowded region. The peripheral of the crowded cells were sparse regions.

### Primary hepatocytes imaging

Mice were euthanized with isoflurane and 25G needle was inserted into the inferior vena cava. After their portal vein is cut, peristaltic pump was used to perfuse the liver. First, Earle’s balanced salt solution (EBSS) (14155063; Invitrogen) containing 5 mM EGTA was used. Following EBSS perfusion, Hank’s balanced salt solution (HBSS) (24020117; Invitrogen) containing 100 U/ml collagenase and trypsin inhibitor was used. The liver was then harvested and massaged to obtain dissociated cells in hepatocyte wash medium (17704024; Invitrogen). After cells went through mesh, the cells were sub-layered onto Percoll (P4937; Sigma). Only viable cells at the bottom layer was washed twice with wash medium, then the cells were plated onto dishes in Wiliams E medium (12551; Invitrogen) containing insulin-transferrin-selenium supplementation. Then the cells were kept in DMEM overnight. Transfection, staining and confocal imaging was performed following the same procedure for HeLa cells.

## Data Availability

All relevant data are available from the authors.

## Acknowledgements

The research was supported in part by the National Institutes of Health (R01-GM115622 and R01-CA207701 to J.W., R01-AG045183, R01-AT009050, R21-EB022302, and DP1-DK113644 to M.C.W., R01-GM120033 to M.M-S.), the Welch Foundation (Q-1912 to M.C.W.), Cancer Prevention and Research Institute of Texas (RP130573 to M.M.S.), the IDDRC Microscopy Core (P30HD024064 and 1U54 HD083092 Intellectual and Developmental Disabilities Research Grant from the Eunice Kennedy Shriver National Institute of Child Health and Human Development), the Optical Imaging and Vital Microscopy core, and the Cytometry and Cell Sorting Core at Baylor College of Medicine with funding from the NIH (AI036211, CA125123, and RR024574) and the expert assistance of Joel M. Sederstrom.

## Author Contributions

X.J., J.C., X.C., M.M-S., T.P., D.M., M.C.W., and J.W. designed experiments; X.J., J.C., C.Z., S.C., Y.Z., M.Z., and X.S. performed experiments; X.J., M.C.W., and J.W. performed data analysis and wrote the manuscript.

